# Identification of Novel Biomarkers for Alzheimer’s Disease and Related Dementias Using Unbiased Plasma Proteomics

**DOI:** 10.1101/2024.01.05.574446

**Authors:** Benjamin Lacar, Shadi Ferdosi, Amir Alavi, Alexey Stukalov, Guhan R. Venkataraman, Matthijs de Geus, Hiroko Dodge, Chao-Yi Wu, Pia Kivisakk, Sudeshna Das, Harendra Guturu, Brad Hyman, Serafim Batzoglou, Steven E. Arnold, Asim Siddiqui

**Author notes:** Correspondence: Asim Siddiqui and Steven E. Arnold. These authors contributed equally (Benjamin Lacar, Shadi Ferdosi, Amir Alavi). Abbreviations: Alzheimer’s disease (AD) and related dementias (ADRD), nanoparticle (NP), mass spectrometry (MS), liquid chromatography mass spectrometry (LC-MS), data-independent acquisition (DIA).

## Abstract

Alzheimer’s disease (AD) and related dementias (ADRD) is a complex disease with multiple pathophysiological drivers that determine clinical symptomology and disease progression. These diseases develop insidiously over time, through many pathways and disease mechanisms and continue to have a huge societal impact for affected individuals and their families. While emerging blood-based biomarkers, such as plasma p-tau181 and p-tau217, accurately detect Alzheimer neuropthology and are associated with faster cognitive decline, the full extension of plasma proteomic changes in ADRD remains unknown. Earlier detection and better classification of the different subtypes may provide opportunities for earlier, more targeted interventions, and perhaps a higher likelihood of successful therapeutic development. In this study, we aim to leverage unbiased mass spectrometry proteomics to identify novel, blood-based biomarkers associated with cognitive decline. 1,786 plasma samples from 1,005 patients were collected over 12 years from partcipants in the Massachusetts Alzheimer’s Disease Research Center Longitudinal Cohort Study. Patient metadata includes demographics, final diagnoses, and clinical dementia rating (CDR) scores taken concurrently. The Proteograph^TM^ Product Suite (Seer, Inc.) and liquid-chromatography mass-spectrometry (LC-MS) analysis were used to process the plasma samples in this cohort and generate unbiased proteomics data. Data-independent acquisition (DIA) mass spectrometry results yielded 36,259 peptides and 4,007 protein groups. Linear mixed effects models revealed 138 differentially abundant proteins between AD and healthy controls. Machine learning classification models for AD diagnosis identified potential candidate biomarkers including MBP, BGLAP, and APoD. Cox regression models were created to determine the association of proteins with disease progression and suggest CLNS1A, CRISPLD2, and GOLPH3 as targets of further investigation as potential biomarkers. The Proteograph workflow provided deep, unbiased coverage of the plasma proteome at a speed that enabled a cohort study of almost 1,800 samples, which is the largest, deep, unbiased proteomics study of ADRD conducted to date.

## Introduction

Dementia affects over 55 million people worldwide, with Alzheimer’s Disease (AD) and Related Dementias (ADRD) being the most common forms. However, heterogeneity in presentation and rates of cognitive decline and disease progression, as well as the need for more informative and accessible biomarkers, contribute to challenges in diagnosis and prognosis. The current gold standard for diagnosis remains autopsy, but this, of course, is only of retrospective clinical and research value. Molecular positron emission tomography imaging (PET, for amyloid-β and tau) approaches the diagnostic accuracy of autopsy, but they are not widely available, accessible or easily repeatable and are expensive. Blood-based biomarkers enable greater accessibility, easier repeatability, and patient participation, ultimately resulting in higher-quality research, disease management, and treatments. Blood-based biomarkers of amyloid beta and phosphorylated tau are emerging with steadily improving accuracy to predict brain AD pathology, but their utility for disease staging or prognosis is still limited. Although AD and related diseases are pathologically defined by their signature proteinopathies, a host of other pathophysiological processes contribute to neurodegeneration and cognitive decline. These include varying degrees of inflammation, vascular disease, metabolic dysfunction, oxidative stress, dysregulation in transcription/translation/post-translational modification, dysproteostasis and dyslipidoses. Much of the heterogeneity of ADRD’s presentations, diagnosis, and prognosis may be related to these factors.

Though liquid chromtagraphy mass spectrometry (LC-MS) remains the gold standard for deep, unbiased proteomics, conducting these experiments in plasma at a scale necessary for biological insight has historically been challenging. Prior studies have been either deep and of limited scale^11^ or at scale but of limited depth.^2^ We previously introduced Proteograph, a platform for deep, unbiased proteomics at scale. Here, we present an updated assay, termed Proteograph XT, to reduce the number of MS injections, enabling a 2.5x improvement in throughput, while preserving similar depth from the Proteograph presented in previous unbiased proteomic studies using nanoparticle-based mass spectroscopy.^3–5^

We therefore used Proteograph XT on 1,786 samples from 1,005 participants, whose final diagnoses represented a spectrum of dementias, with AD participants (n=379) representing the plurality. With these data, our study addressed biological pathways that are implicated in AD using linear mixed modeling and differential expression, biomarker discovery for AD patients with machine learning classification, and potential biomarkers for cognitive decline across dementia patients with time-to-event modeling.

## Methods

### Cohort

The cohort consisted of 1030 participants in the Massachusetts Alzheimer’s Disease Research Center’s Longitudinal Cohort Study (MADRC-LC) in whom at least one plasma had been collected between 2008 and 2019. This is a longitudinal observational study spanning the entire continnum of nomal aging to ADRD. Annual standardized assessments included a general and neurological exam, a semi-structured interview with the participant and/or informant to record cognitive symptoms with a Clinical Dementia Rating scale (CDR Dementia Staging Instrument), a battery of neuropsychological tests, and other instruments of the National Alzheimer’s Coordinating Center (NACC) Uniform Dataset (UDS)^6,7^. Blood was collected from all consenting participants.

Cognitive status and clinical syndromes were determined at each visit by a consensus team after a detailed examination and review of all available information according to the 2011 NIA-AA diagnostic criteria for MCI and AD^8,9^. Many participants had autopsy, imaging, CSF, and/or plasma biomarkers in affiliated protocols. Disease diagnosis (AD or other diseases) was further informed by these data when available. Participant clinical data used in the analyses here include age, sex, race, ethnicity, years of education, and clinical dementia rating global (CDRg) scores taken concurrently with sample collection. Additional biomarker data available on almost all cases included apolipoprotein e (APOE) genotype as well as plasma phospho-tau 181 (pTau181), glial fibrillary acidic protein (GFAP) and neurofilament -light (NfL). Plasma biomarkers were measured using ultrasensitive MSD S-PLEX electrochemiluminescence immunoassay kits (Meso Scale Discovery, Rockville, MD), as previously described^1010^.

Participant summary statistics are shown in Table 1.

**Table 1.** Study participant summary statistics.

| Characteristics | Alzheimer's Disease | No Neurodegenerative Disease | Other Dementias |
| --- | --- | --- | --- |
| Number of participants (N) | 379 | 240 | 387 |
| Number of visits [median (min, max)] | 1 (1, 6) | 2 (1, 6) | 1 (1, 6) |
| Age at 1<sup>st</sup> visit [Mean (SD)] | 74.5 (9.4) | 66.7 (11.6) | 71.4 (9.9) |
| Female (%) | 190 (50.1) | 166 (69.2) | 212 (54.8) |
| College educated [N (%)] | 264 (69.7) | 165 (68.8) | 253 (65.4) |
| APOE ε4 carriers [N (%)] | 185 (48.4) | 55 (22.9) | 82 (21.2) |
| Last Draw CDR sum of boxes [Mean (SD)] | 5.0 (4.5) | 0.2 (0.5) | 3.9 (4.6) |
| Last Draw MMSE [Mean (SD)]<sup>1</sup> | 18.0 (8.2) | 26.6 (2.8) | 21.5 (7.2) |
| Last Draw pTau-181 (pg/mL) [Mean (SD)]<sup>1</sup> | 4.7 (3.3) | 2.5 (1.3) | 2.5 (1.4) |
| Last Draw NfL (pg/mL) [Mean (SD)]<sup>1</sup> | 319.4 (213.0) | 180.2 (106.7) | 326.7 (322.9) |
| Last Draw GFAP (pg/mL) [Mean (SD)]<sup>1</sup> | 146.9 (125.5) | 80.5 (44.2) | 99.2 (66.1) |

Standard Protocol Approvals, Registrations, and Patient Consents

The study was approved by the Mass General Brigham Institutional Review Board (2006P002104) and all participants or their assigned surrogate decision makers provided written informed consent.

### Sample Preparation

Plasma samples used in this study had been collected between 2008 and 2019 and were banked in the Harvard Biomarkers Study Biobank^11^. Samples were collected in K2EDTA tubes, centrifuged at 2000 g or 5 min, frozen in low retention polypropylene cryovials within 4 hours of collection and stored at -80°C until use.

Plasma from 1,786 individual samples (including subsequent plasma collection samples from the same individuals) and a plasma control sample (PC6), consisting of pooled citrate phosphate dextrose anticoagulant plasma from 15 healthy individuals, were processed with the Proteograph XT Assay Kit (Seer). Plasma tubes containing 240 µL of plasma were loaded onto the SP100 Automation Instrument (Seer) for sample preparation to generate purified peptides for LC-MS analysis. The samples were incubated to form each of the two proprietary, physicochemically distinct nanoparticle (NP) suspensions for protein corona formation. Samples (40 samples/plate; 38-39 individual plasma samples and 1-2 PC6 samples) were automatically plated, including process controls, digestion control, and MPE peptide clean-up control. After a one-hour incubation, leveraging the paramagnetic property of NPs, NP-bound proteins were captured using magnetic isolation. A series of gentle washes removed nonspecific and weakly bound proteins. This process results in a highly specific and reproducible protein corona. Protein coronas are denatured, reduced, alkylated, and digested with Trypsin/Lys-C to generate tryptic peptides for LC-MS analysis. All steps were performed in a one-pot reaction directly on the NPs. The in-solution digestion mixture was then desalted and all detergents were removed using a solid phase extraction and positive pressure (Monitored Multi-Flow Positive Pressure Evaporative Extraction module [MPE]^2^ ^TM^; Hamilton) system on SP100 Automation Instrument. Clean peptides were eluted in a high-organic buffer within a deep-well collection plate and quantified. Equal volumes of peptide elution were dried down in a SpeedVac (3 hours-overnight), and the resulting dried peptides were either reconstuited for immediate analysis by liquid-chromatography mass-spectroscopy (LC-MS) or stored at -80 °C to be analyzed later. Peptides were reconstituted to a final concentration of 0.06 µg/µL in Proteograph XT Assay Kit Reconstitution Buffer.

### LC-MS Analysis

8 µL of the reconstituted peptides were loaded on an Acclaim PepMap 100 C18 (0.3 mm ID x 5 mm) trap column and then separated on an Ultimate 3000 HPLC System and a 50 cm μPAC HPLC column (Thermo Fisher Scientific) at a flow rate of 1 μL/minute using a gradient of 5 – 25% solvent B (0.1% FA, 100 % ACN) in solvent A (0.1% FA, 100% water) over 22 minutes, resulting in a 33-minute total run time. For the MS analysis on the Thermo Fisher Scientific Orbitrap Exploris 480 MS, 480 ng of material per NP was analyzed in DIA mode using 10 m/z isolation windows from 380-1000 m/z. MS1 scans were acquired at 60k resolution and MS2 at 30k resolution.

### Spectral Library Generation

#### Gas Phase Fractionation (GPF)

We used an MS-only workflow that combines GPF and DIA LC-MS, saving significant experiment time while maintaining high data completeness and reproducibility^1^. This strategy generated a chromatogram spectral library with GPF deep scanning experiments, consisting of staggered m/z window analysis of the pooled peptides left over from Proteograph XT Assay plates by pooling up to 5 µL of tryptic peptides left for each sample in the plate into separate pools for each NP suspension. Six DIA LC-MS injections of 10 µL each containing a peptide concentration of 0.06 ug/µL from each NP pool were analyzed. The six injections covered mass over charge (m/z) ranges of 400-500 m/z, 500-600 m/z, 600-700 m/z, 700-800 m/z, 800-900 m/z, and 900-1000 m/z, with each injection having 50 staggered windows covering 4 m/z. MS1 was run in 60K resolution and MS2 was run in 30K resolution on another Orbitrap Exploris 480 MS with similar chromatographic setup (LC, trap, and column). A library-free search of the DIA LC-MS data was performed using DIA-NN 1.8.1^12^ to create the empirically corrected GPF library.

### Data Analysis and Protein Representation

All MS files were developed to run DIA-NN 1.8.1 with a GPF library search. All identifications are reported at 1% FDR. Panel protein representations integrated nanoparticle:precursor representations with MaxLFQ^13,14^.

### Functional Annotation Enrichment and Differential Expression Analysis

To determine how the biological variables in this cohort correlate with protein abundances comparing profiles across the 4,007 protein groups and 1,786 plasma samples, we trained a linear mixed-effects model (LMM; *lme4*) with ProteinIntensity ∼ Diagnosis + Diagnosis:(Age + Sex + Education + globalCDR + ApoE_score) + Education + Sex + Age + SampleVariation + (1|CollectionYear) + (1|NP:AssayPlate), where Diagnosis contains 3 levels of AD, other dementia, and no neurodegenerative disease, and ApoE score is calculated as (-0.5 * n of e2 alleles + 1 * n of e4 alleles). SampleVariation is a technical variable that accounts for variabilities in the samples resulting from differences in NP:protein interactions that are due to variations in sample collection. We calculated this variable the median fold-change of proteins annotated as “Nucleolus” for each plasma sample and NP. We picked Nucleolus as the term describing the sample variation here because we observed the highest variation between samples with this term compared to other GOCC terms such as “extracellular”, “intracellular”, “cytoskeleton”, and “humoral immune response”. CollectionYear is included as a random effect to account for sample variability based on the year of sample collection, and NP and assay plates associated with the NP is accounting for sample preparation variabilities. The missing protein intensities are imputed for the NP that has the lowest number of missingness across all samples, and in the case of equal missingness the NP with higher protein intensity is picked for imputation. The imputation is done by sampling 3 times from a shifted normal distribution for that feature with mean shift = -1.8 and width = 0.3^15^.

To determine functional annotations associated with the LMM results, annotations were matched with UniProt identifiers and enrichments calculated based on the coefficient distributions using the R AnnoCrawler package and implementation of 1D annotation enrichment^16^.

To indicate how proteins are differentially abundant in AD cases in contrast to the group without neurodegenerative disease, the LMM coefficients, where Diagnosis = AD, were plotted against the negative log10 transformed p-values where the p-values are corrected for multiple testing according to the Benjamini-Hochberg method (Figure 2B).

**Figure 1.**
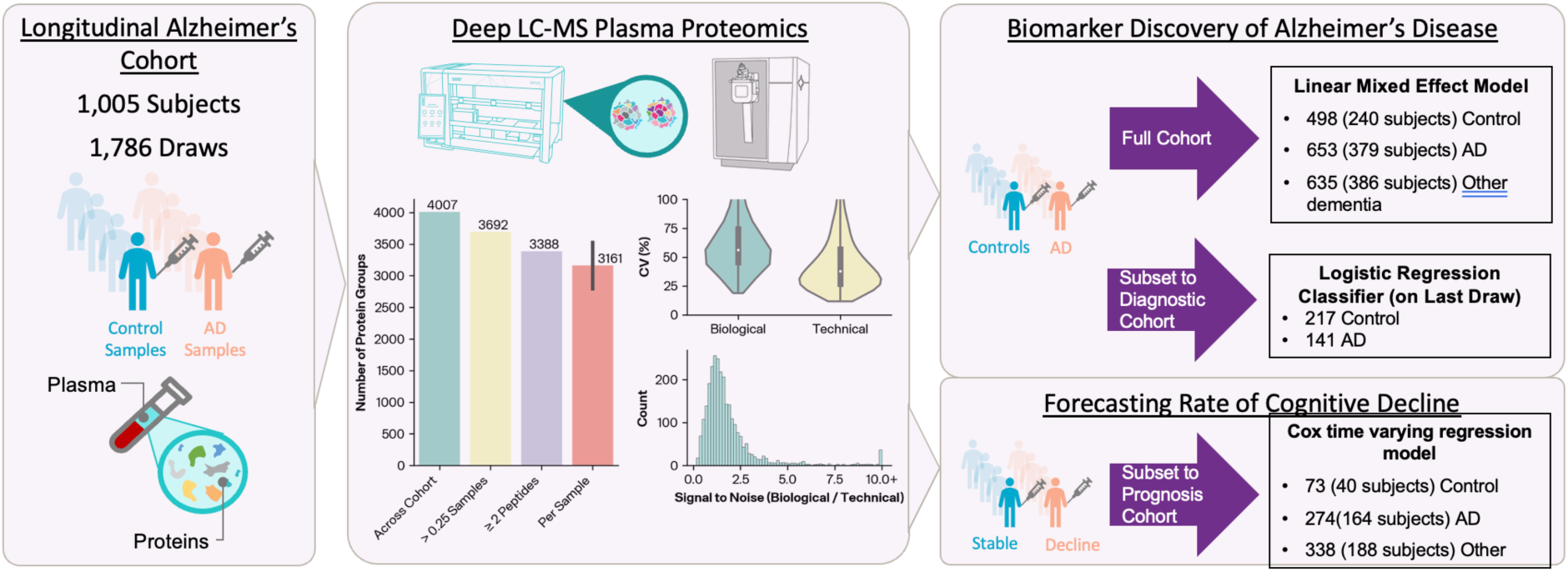
Study design, plasma proteomics quantifications, and cohort subsetting for questions of interest.

**Figure 2.**
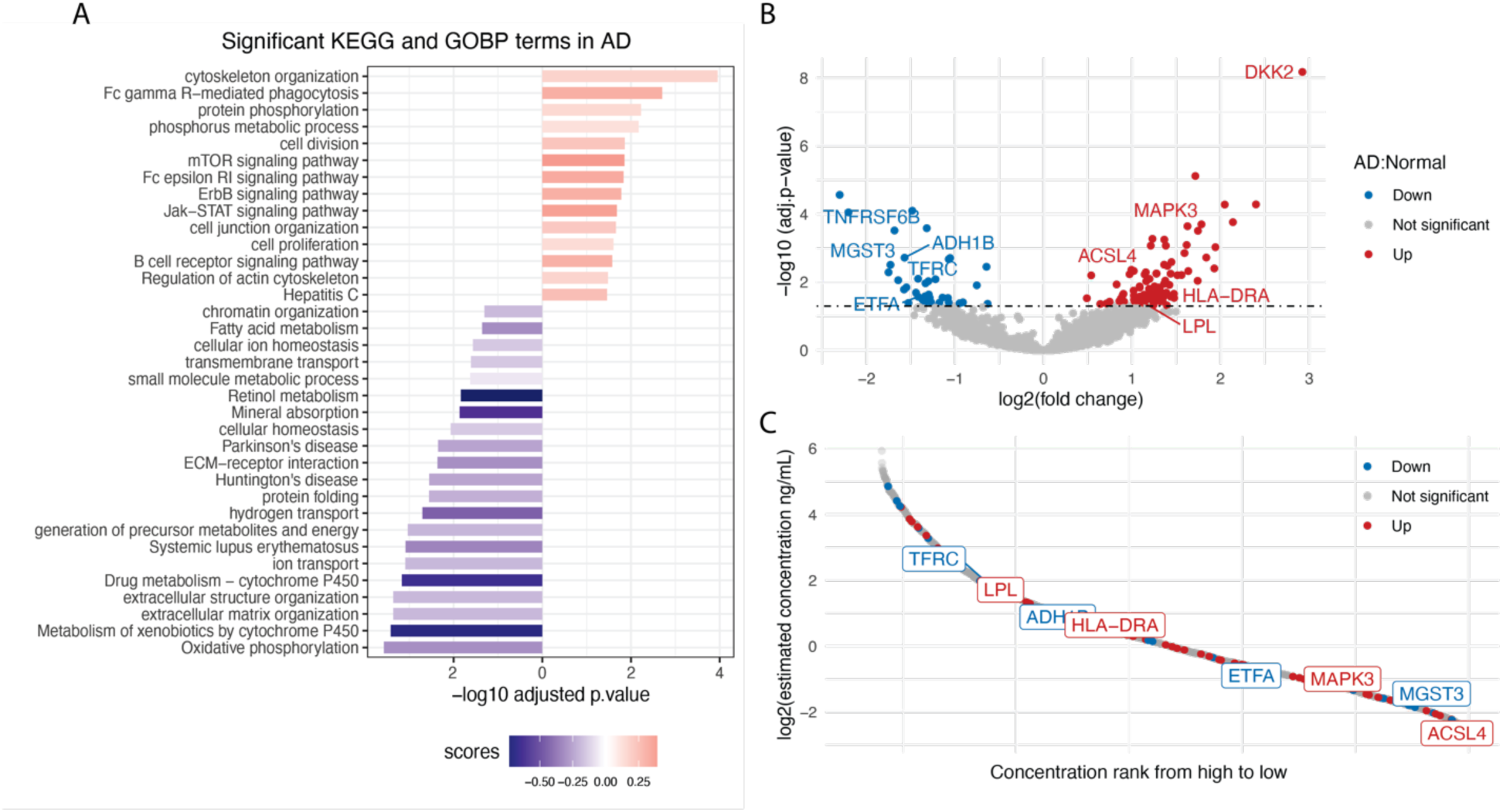
Pathway enrichment analysis and differential expression analysis in Alzheimer’s disease. A) 1D annotation enrichment analysis for AD. 1D score was calculated for KEGG and GOBP terms as described in Method. The p-values are adjusted based on Benjamini-Hochberg (BH) multiple testing correction and filtered at 5% FDR. Enrichment (score > 0) is depicted in red; Depletion (score < 0) is depicted in blue. B) Differential expression analysis for AD vs. healthy controls. Volcano plot showing log2 fold change of proteins (x axis) and -log10 p-values after BH multiple testing correction (y axis). The upregulated proteins are depicted in red; the downregulated proteins are depicted in blue. C) Dynamic range of identified proteins matched with HPPP database^37^. All proteins identified in the cohort that could match to the HPPP database shown in grey; upregulated protein in AD shown in red; downregulated proteins in AD shown in blue.

### Diagnostic Cohort and Machine Learning Diagnostic Model

We established a Diagnostic cohort using the final draw from each sample for the purpose of evaluating the use of protein biomarkers for determining AD status and identifying AD related proteins. We then further restricted these data to only those participants which diagnosed as “AD” or “No Neurodegenerative Disease”. As we wished to evaluate pTau-181 as a biomarker and to avoid confounding of diagnostic state, we also excluded cases where the diagnosis was made on the basis of pTau-181. Ideally, we would remove all cases which used a biomarker to determine AD status, but this would yield too few healthy controls. The final set of samples included 141 AD participants, and 217 healthy controls.

We developed a machine learning model to classify AD and Healthy controls based on their plasma proteomics features from LC-MS, in addition to pTau-181 concentration. Our logistic regression model includes a preprocessing pipeline for the proteomics features that appropriately handles missing data, imputation, normalization, and feature selection (Figure 3a). Protein intensities are first filtered by missingness, keeping only features that have a missing rate of less than or equal to 75% among the training samples. We then normalize the features by taking the logarithm and subtracting each feature’s median. Any remaining missing values are imputed by sampling from a shifted normal distribution for that feature with mean shift = -1.8 and width = 0.3^1515^. The top-K features are then identified by computing the ANOVA F-score between the labels and features, and keeping the K highest scoring features. Finally, pTau-181 (pg/mL) is added as a feature and the whole set of features is mean centered and unit variance scaled, before passing to a penalized logistic regression classifier.

**Figure 3.**
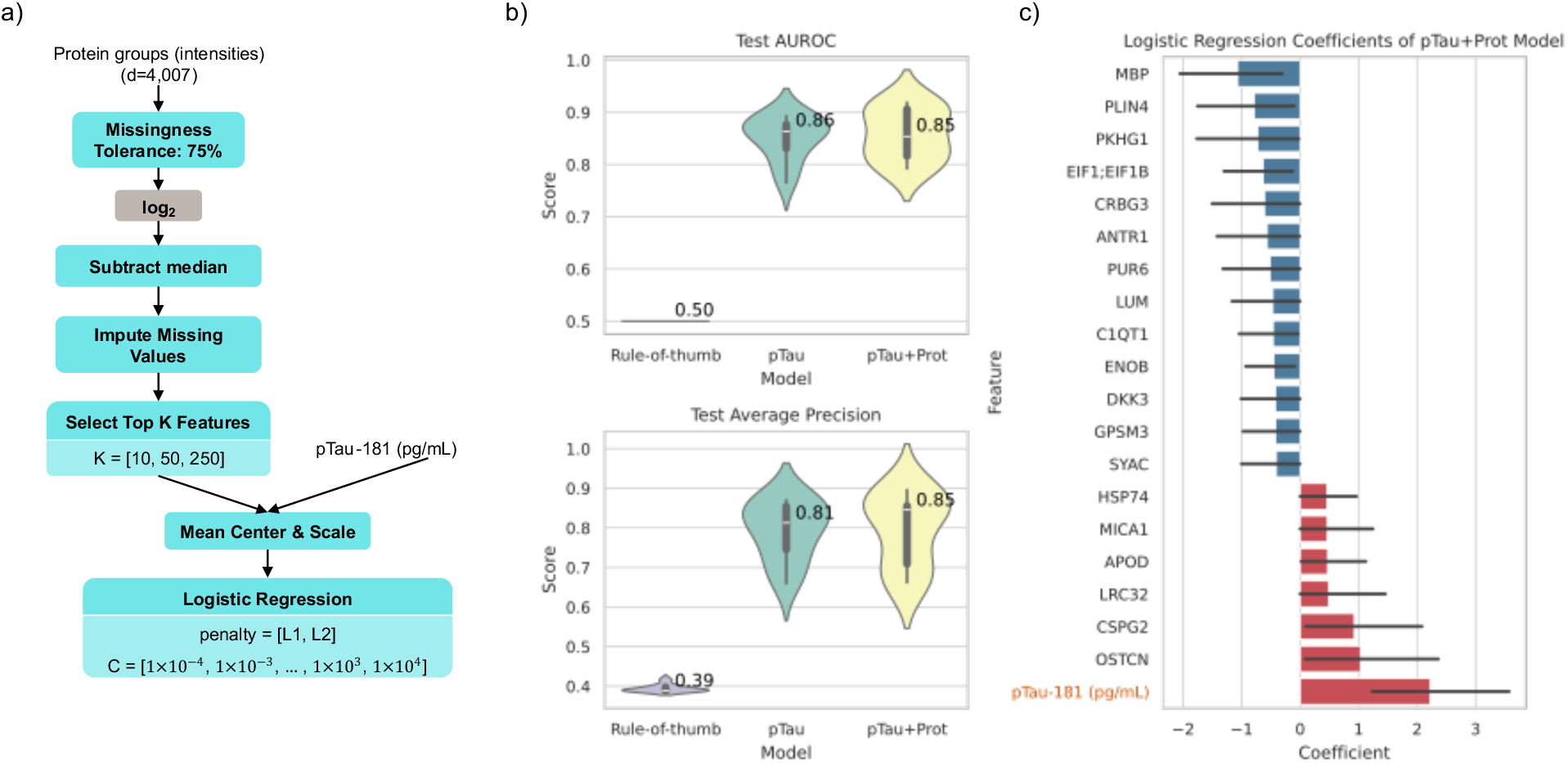
Classification of AD vs Healthy controls. A) Flowchart of our machine learning pipeline for our “pTau+Prot” model. Blue nodes are steps that fit parameters based on training data and apply them to validation and test data (e.g. a list of features that pass missingness filters, the median value for a feature, or the coefficients of the logistic regression classifier. Grey nodes are parameterless. Some nodes have hyperparameters listed, which we tune using nested cross validation. B) Results of 10 fold cross validation comparing the AUROC and Average Precision of the Rule-of-thumb baseline classifier, pTau-181 concentration alone, and our model shown in panel (A). C) The top 20 average coefficient values for the pTau+Prot model (when sorted by absolute values) across the 10 folds. Error bars indicate 95% confidence internal estimated using 1000 bootstraps with replacement.

The model described above has hyperparameters (K for feature selection, and penalty kind and amount for logistic regression) that must be tuned, and logistic regression coefficients that must be fit to the data. To avoid overfitting, we adopt a nested cross-validation strategy. We create 10 outer folds. For each of these 10 folds, the other 9 are taken as the training set. This training set is then further split in an inner hyperparameter tuning stage, where 80% of it is used to fit a model for each possible hyperparameter setting, and the other 20% is a validation set used to evaluate the hyperparameter setting. The best hyperparameter setting (highest area under the receiver operator characteristic curve (AUROC)) is then refit on the full 9-fold training set, and a test score is computed on the test fold.

### Time-to-Event Analysis

Cox proportional hazards (CPH) and Cox time-varying (CTV) regression models were built to determine the association of each protein group with the time to CDRg increase (the event) from either CDRg of 0.0 or CDRg of 0.5. Participants who showed an increase from baseline (0.0 to 0.5 or 0.5 to 1) after a minimum of 1 post-draw visit were labeled as E=1 while those that did not show an increase for their observation time and for at least 3 years were categorized as E=0 (censored). With these criteria, the original dataset was subset in the CDRg baseline 0.0 model with 300 participants and 540 biosamples (n=145 participants with multiple draws) and the CDRg baseline of 0.5 model had 391 participants and 684 biosamples (n=209 participants with multiple draws). 70 participants were in the models for both baselines. We describe three sets of Cox regression model types in this study:

**Table 2.**

|  | Model type | Patient samples | Uses Delayed Entry | Timescale | Purpose |
| --- | --- | --- | --- | --- | --- |
| Model 1 | CTV | All before event | Yes | Age | Represent multiple blood draws and time-varying protein levels |
| Model 2 | CPH | Last before event | Yes | Age | Comparison with CTV model |
| Model 3 | CPH | Last before event | No | Follow-up time | Check of PH assumption and generation of survival curves |

The equations for these models are shown below, with a detailed explanation of predictor variables to follow:

Model 1: *h*(*a*, *X*(*t*)) = *h*_0_(*a*|*a*_0_) *exp*[Σβ_*δ*_X_i_ + Σδ_j_X_j_(*t*)5

Model 2: *h*(*a*, *X*) = *h*_0_(*a*|*a*_0_) *exp*[Σβ_*δ*_X_i_]

Model 3: *h*(*t*, *X*) = *h*_0_(*t*) *exp*[Σβ_*δ*_*X*_*δ*_]

The variables common to all three models are *h* representing the hazard rate, β_*δ*_, the regression coefficient corresponding to X_i_, and *X*, a vector of time-independent covariates. In Model 1 only, δ_j_represents the coefficients of X_j_ and *X*(*t*) is a vector of time-dependent covariates. In Models 1 and 2, *a* is the age at event or censorship, *a*_0_is the age of last blood draw before the event or censorship, indicating that both models adjust for delayed entry and use age as time-scale. In Model 3, the variable *t* represents the duration between the last blood draw and the event or censorship, does not account for delayed entry, and instead age of the last blood draw is used as a covariate.

The time-varying model (Model 1) maximizes the data available from participants with multiple blood draws and represents proteins as a time-varying covariate. Models 2 and 3 only used the last available blood draw before an event. The last draw model showed a greater correlation with the CTV model than the first draw model (Supp. Fig X) and therefore the last draw was the basis for assessing the proportional hazards assumption and survival curve generation in Model 3. Models 1 and 2 used age as time-scale given the importance of age in dementia^17^. Both models also accounted for delayed entry, where entry time is the age of earliest draw at a subject’s baseline since the MADRC cohort is an observational study with an open cohort^18^. Model 3 did not use age as time-scale and used age as a covariate instead. All models assessed the association of each protein group while controlling for the subject-level covariates sex, education, ApoEe4 risk score (-0.5 * n of e2 alleles + 1 * n of e4 alleles) and technical-level covariates that contributed to variation of the protein group itself (Supp. Figure X), including plate group identifier, collection year, and nucleolus score. Models were created for a protein group only if there was a minimum completeness of 25%; samples with missing values for a protein group were not considered in that model. The intensity values of each protein group were median normalized, log2 transformed and standardized. Since one model was built for each protein group, we accounted for multiple hypothesis testing by applying Benjamini-Hochberg adjustment to nominal p-values. In partial regression coefficient plots, the levels of protein group features are shown as z-scores. The python package *lifelines* was used to create the Cox models^19^.

## Results & Discussion

### Study design and protein quantification metrics

Samples were collected from an observational study of a group of individuals with or without cognitive impairment in a longitudinal fashion with data collected on a nearly annual basis. Data include cognitive tests and blood collection (average 6.2 ± standard deviation 3.80 visits per subject), although proteomics was obtained for only a subset of blood draws (1.8 ± 1.04 blood draws per subject). Final primary disease diagnoses were also provided along with a method of determination such as neuropathology, molecular neuroimaging, CSF and/or plasma biomarker. Plasma samples were processed for deep LC-MS proteomics. In our analysis of 1,786 plasma samples, we identified 4,007 protein groups (3,692 for those in at least 25% of samples) and 36,259 peptides using the GPF library.

### Differential Expression of Proteins in Alzheimer’s Patients

To investigate the biological pathways involved in AD and the identification of proteins that are differentially abundant between AD and control samples, we analyzed all 1786 plasma samples, including 498 plasma samples from participants without neurodegenerative disease, 653 plasma samples from participants with AD, and 635 plasma samples from participants with other types of dementia.

We use a linear mixed model describing the normalized intensities of all identified proteins as a function of diagnosis, age, sex, education, global CDR score, APOE alleles, and technical variables such as NPs, assay plates, sample collection year, and plasma protein composition. The resulting coefficients from this model were then used in a 1D annotation enrichment analysis to evaluate how these biological variables are differentially associated with functional annotations. Figure 2A shows the biological processes that are significantly dysregulated in AD. For example, oxidative phosphorylation is shown to be downregulated (Enrichment score = -0.34). In a recent study by Misrani et. al, mitochondrial dysfunction, associated with a decrease in neuronal ATP levels, has been shown to be a characteristic feature of AD. This dysfunction is partly due to the overproduction of reactive oxygen species (ROS), leading to oxidative stress and damage to mitochondrial function. In AD, this results in compromised oxidative phosphorylation, leading to neuronal cell death^20^. The extracellular matrix (ECM) is another biological processes that has been shown to be dysregulated in AD (Figure 2A). Dysregulation of ECM plays a significant role in its pathogenesis, and it is involved in various aspects of AD, including synaptic transmission, amyloid-β plague generation and degeneration, tau-protein production, oxidative stress response, and inflammatory response. Alterations in ECM components can affect the stability of perineuronal nets, impacting the clearance of amyloid-β and the production of neurofibrillary tangles^21,22^. Signaling pathways such as mTOR, ErbB, and Jak-STAT that are shown to be upregulated in AD participants in this dataset, are known pathways related to the pathogenesis of AD^23–25^ (Figure 2A). In addition, pathways related to Parkinson’s disease (PD) and Huntington’s disease (HD) are shown to be significantly dysregulated. Although each of these diseases has its unique pathophysiological mechanisms, they do share some common mechanisms, including misfolding and aggregation of beta-amyloid and α-synuclein, leading to neuronal apoptosis^26,27^.

To gain insights into which proteins are differentially abundant in the plasma of AD patients compared the control group, we performed a differential expression analysis using the same linear mixed model as that above. This analysis resulted in 138 differentially abundant proteins of which 38 are down-regulated proteins and 100 up-regulated proteins (Figure 2B). For instance, MAPK3, one of the up-regulated proteins in AD participants in this dataset, is known to play a crucial role in AD^28^. The MAPK pathways, including extracellular signal-regulated kinase (ERK), c-Jun N-terminal kinase (JNK), and p38 pathways, are activated in neurons vulnerable to AD. The MAPK pathways are linked to significant pathological processes in AD, such as tau phosphorylation, amyloid-beta deposition, and amyloid-beta protein precursor functioning^29^. ACLS4 is another upregulated protein in AD participants. ACLS4 is involved in the regulation of synaptic function and neuronal signaling and previous studies have shown that its level is significantly increased in AD patients^30^. DKK2 which is the most up-regulated amongst AD participants in this dataset, is an inhibitor of the Wnt signaling pathway, which is known to be crucial for cognitive function, and its upregulation may contribute to reduced WnT signaling in AD^31–33(Figure^ 2B).

MGST3 which is one of the downregulated proteins in AD participants (Figure 2B), is known to be significantly associated with hippocampus size and found to be linked to neurodegenerative disorders associated with reduced hippocampus volume such as AD, PD, and HD^34^. ADH1B which we found to have a protective association in our data, has also been found to suppress Aβ-induced neuron apoptosis^35^, and mutations in its gene have been found to be associated with the development of AD^36^.

To investigate how abundant these signature proteins are in blood plasma, we mapped the identified proteins in our cohort to the Human Plasma proteome (HPPP) Database^37^. Figure 2C shows that the dysregulated proteins are distributed across the dynamic range of the plasma proteome with some of the highlighted proteins such as MAPK3, MGST3, and ACSL4 being at the lower abundance range. In this regard, our finding of their differential expression in AD is noteworthy because these proteins would have been assayed in a unbiased proteomics assay in a large cohort without the use of Proteograph XT.

### Biomarker-Based Classification of Alzheimer’s Patients

Next, we investigated if there is a multimarker signature of AD that can be identified from the proteomics data. While pTau measurement has been established as the best marker for determining AD status, we were curious to see if protein features could provide additional evidence of an AD signature beyond known autopsy, PET, CSF and plasma biomarkers. We also aimed to use this approach to determine AD related proteins. To this end, we focused on a subset of samples using the last draw from each subject that has at least two clinical visits. We then further restricted these data to only those participants which diagnosed as “AD” or “No Neurodegenerative Disease” and took care to ensure that there was no confounding information through inclusion of cases where the diagnostic status is based on biomarkers we intended to evaluate. The final set of samples included 141 AD participants, and 217 healthy controls.

We developed a logistic regression-based machine learning model to classify AD versus healthy controls using both pTau-181 concentration and our LC-MS proteomics features and evaluate it using nested cross validation (Methods, Diagnostic Cohort and Machine Learning Diagnostic Model). Since our dataset is imbalanced (AD is the minority class), we report the average precision (average positive predictive value) in addition to AUROC (Figure 3b). We compared our model to a rule-of-thumb classifier that would just report the prior class distribution as the predicted probability of AD, in addition to comparing to the value of pTau-181 concentration alone.

We can see in Figure 3b that the model using proteomics features (“pTau+Prot”) does not have a significant increase in diagnostic performance over pTau-181 on its own. Furthermore, we plan to measure plasma pTau-217 levels in these samples which we anticipate will provide better discrimination of AD status compared to our general protein model. We hypothesize that this is because pTau markers have been developed highly studies in relationship to AD leaving little room for improvement and we do not measure phosphorylation state in the unbiased proteomics assay. Nevertheless, we can use these models to give us potential insight into biomarkers driving the various pathophysiological processes that contribute to neurodegeneration and cognitive decline.

To that end, we can interrogate the fitted models to determine which input features were most influential in classifying AD and healthy controls. The average of the logistic regression coefficients across the 10 models (from the 10 folds) was computed, and the top 20 (based on absolute value) are reported in Figure 3c. While pTau-181 concentration was the most influential feature, other noteworthy proteomics features also had large coefficients. Myeloid basic protein (MBP) was associated with Healthy controls (negative coefficient, protective), and prior studies have shown that MBP acts as an amyloid β-protein (Aβ) chaperone and can be an inhibitor of accumulation of Aβ fibrils^38–44^. Other studies have found the opposite association as well, and the relation of MBP to AD pathology is still an open area of research^45^. In addition^45^ to osteocalcin (OSTCN), a marker of processes involved in osteoporosis such as bone remodeling and anabolism, and prior studies have shown some comorbidity of AD and osteoporosis^46^. Apolipoprotein D (APOD) also had a large positive coefficient and is known to have increased levels in AD where it plays a neuroprotective role against oxidative stress^47,48^.

### Cox regression models identify multiple biomarkers associated with dementia progression

To determine the association of protein groups to dementia progression, we employed multiple Cox regression models where we assessed the time to CDRg increase. The primary model (Model 1) is a Cox time-varying model that represents delayed entry, due to open cohort enrollment, right-censored events, and age as timescale. The time-varying component of this model allows the protein expression to be represented over time when multiple draws are available for a subject (Fig. 4a, b) and for the proportional hazard assumption to be relaxed. We reasoned that cognitively healthy controls (CDRg of 0.0) would have different rates of dementia progression than those already showing mild cognitive impairment (MCI, CDRg of 0.5). We therefore built one model for participants with a baseline CDRg=0 and another for baseline CDRg = 0.5. The distribution in participants’ final diagnoses for non-neurodegenerative, AD, and other dementias was 179, 43, and 78 participants respectively in the CDRg 0.0 cohort and 40, 164, and 188 participants, respectively, in the CDRg 0.5 cohort.

**Figure 4.**
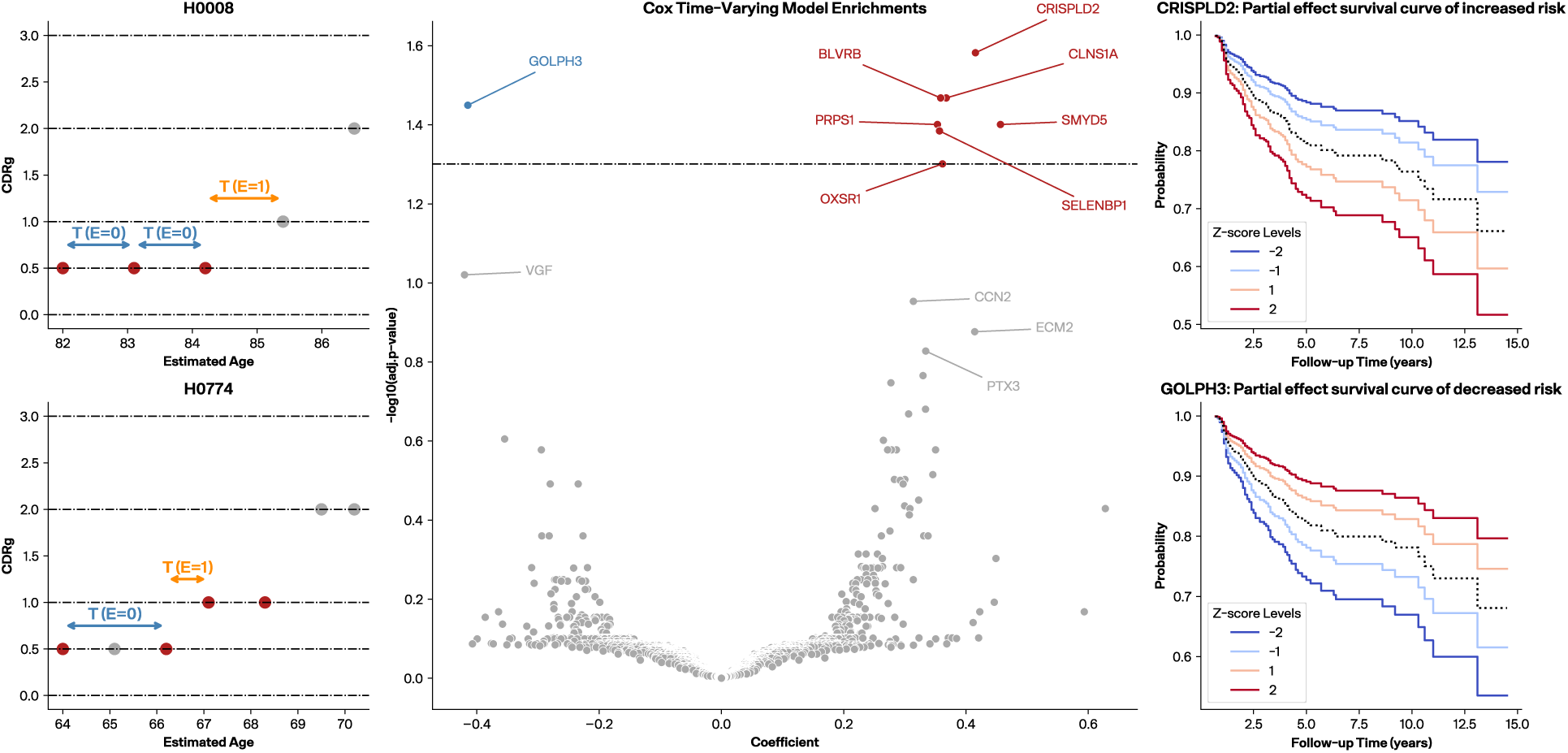
Time-to-CDRg increase assessments with Cox regression models. (a, b) Two examples of participants’ CDR history. All periods represented in a Cox time-varying model between obtained blood samples and before an event are characterized as E=0 and the period between last blood draw and an observed event (CDRg increase) is characterized as E=1. A subject must have at least three visits without a CDRg increase for the final period to be labeled as censored (E=0). (c) CTV volcano plot where the y-axis is -log10 adjusted p-value (BH-corrected) and the dashed line represents adjusted p-value of 0.05. Proteins with positive coefficients are indicative of proteins associated with increased risk of time-to-CDRg increase while those with negative coefficients are associated with decreased risk. (d) CRISPLD2 survival curve generated from a CPH model between last draw and event as an example of a positive coefficient. (e) GOLPH3 survival curve generated from a CPH model between last draw and event, as an example of a negative coefficient.

We found that the CTV models for events greater than 0.0 model had no protein groups significantly associated with time-to-CDRg increase. However, the CTV models for events greater than 0.5 identified eight protein groups with coefficients that were significantly associated (p-adj < 0.05 after BH correction) (Fig. 4b). Seven protein groups had positive coefficients indicating that elevated levels would implicate a shorter time-to-CDRg increase: CRISPLD2 (Q9H0B8), CLNS1A (P54105), BLVRB (P30043), SMYD5 (Q6GMV2), PRPS1 (P60891), SELENBP1 (Q13228_Q13228.4), and OXSR1 (O95747). One significantly associated protein group GOLPH3 (Q9H4A6) had a negative coefficient, implying that higher levels are associated with delays in CDRg increase. Additionally, VGF (O15240), identified as a significant biomarker of AD in brain tissue, CSF and mouse model proteomics studies^49–51^, but not previously in plasma, was just outside significance (p-adj < 0.1 after BH correction) and was also negatively associated with the time-to-CDRg increase.

To further assess our time-to-event approach, we evaluated a CPH model using the latest draw available (nearest to but before the event) with delayed entry and age as timescale (Model 2). We found that the CPH models did not show any significantly associated proteins after BH correction, demonstrating that the CTV model provided greater statistical power than the CPH models (Supp. Fig X.) However, amongst those in the top 20 of lowest nominal p-values of Model 2 were six proteins (CLNS1A, CRISPLD2, GOLPH3, OXSR1, PRPS1, SELENBP1) that were also significantly associated in the CTV model. A different CPH model, one without delayed entry and age of the blood draw as a covariate (Model 3), was used to assess the proportional hazard assumption and generate survival curves, with positive and negative association examples with time-to-CDRg increase shown in (Fig 4 d,e).

Several proteins identified in the CTV and CPH models showed relevance to dementia and/or Alzheimer’s disease in prior studies. CRISPLD2 was identified as one of 89 genes regulated in an AD blood transcriptome study that accounted for white matter hyperintensities^52^. CLNS1A was one of the significant variably methylated probes associated with amyloid-β in postmortem dorsolateral prefrontal cortex^53^. GOLPH3, which promotes vesicle exit for trafficking to the plasma membrane, has not been implicated directly in dementia or AD, but it was cited as a potential mechanism for Golgi fragmentation in AD^54^. Studies on OXSR1^55^, PRPS1^56^, and SELENBP1^57^ also show indirect evidence for these proteins in dementia. In addition, a number of proteins just above the BH cutoff of 0.05 have greater support for a role in dementia/AD, including VGF, MMP9, and CCN2.

## Discussion

The goal of this study were to leverage deep, unbiased plasma proteomics to identify biomarkers associated with dementia progression and Alzheimer’s disease. We employed a variety of approaches using different subsets of the proteomics data to discover biological pathways relevant to Alzheimer’s disease, uncover biomarkers of disease classification, and reveal proteomic signatures of dementia progression. While some of the pathways and proteins identified are known to be involved with ADRD, many are not and may point to novel biology. The proteins associated with dementia progression are of particular interest. They may enable development of a model to predict individuals that are at risk of rapid cognitive decline. Such a model could be used to aid treatment decisions in patients.

Some strengths of the study include the large sample sizes, the standardized clinical characterization of cognition and function over time, the depth of the plasma proteome covered that is enabled by Seer’s Proteograph workflow and GPF and DIA LC-MS workflow, and the Cox regression models to identify those proteins most associated with clinical prognosis.

The open cohort and volunteer enrollment of the study implicates a bias in observed time for participants compared to a randomized controlled trial. In addition, participants may preferentially enroll if they or their caregivers notice signs of dementia, as observed in the final diagnoses of the CDRg 0.5 cohort. Nevertheless, we aimed to minimize these sources of bias by using Cox models with appropriate modeling parameters including delayed entry, age as time-scale, right censorship, and time-varying protein covariates. Our cohort was predominantly composed of people of white race, European ethnicities and high education, thus limiting our ability to generalize findings to people of non-European ancestry and less education who are under-represented in AD research.

A major advance of this work is the use of the next iteration of the Proteograph platform for deep, unbiased proteomics. This platform allowed us to run a large study of almost 1,800 samples while assaying over 4,000 proteins and 36,000 peptides. This depth at this scale was not previously possible for an unbiased workflow. As reported elsewhere, with newer MS analyzers this workflow can achieve 6,000 proteins and more. Unbiased discovery provides an opportunity to learn new biology and develop a deeper understanding of disease. It also provides an opportunity for peptide and hence isoform level analysis. Future work could investigate those aspects of the data in more detail as well as attempting to dissect the similarities and differences in the pathophysiological pathways associated among ADRD, as well as heterogeneities among AD stage and subtypes.

## Acknowledgments

We thank Daniel Hornburg and Khatereh Motamedchaboki for their comments and feedback on the project. We thanks the Seer Operations team for processing the plasma samples.

This work was funded in part by SBIR Award 5R44AG065051-02 granted by the National Institute of Aging, part of the National Instistute of Health.

## Footnotes

1 Not available in all participants, statistics based on non-null values.

